# Large-Scale Structure-Based Virtual Screening Identifies Diverse K_Na_1.1 (KCNT1) Potassium Channel Inhibitors

**DOI:** 10.1101/2025.09.30.679465

**Authors:** Emily A. Caseley, Katie J. Simmons, Bethan A. Cole, Alex J. Flynn, Stephen P. Muench, Jonathan D. Lippiat

## Abstract

Severe drug-resistant childhood epilepsy is caused by *KCNT1* gain-of-function genetic variants, resulting in increased K_Na_1.1 channel activity. *KCNT1*-associated epilepsy is thought to affect around 1 in 300,000 births worldwide. Current treatment for KCNT1 epilepsy only provides mild symptomatic relief and uses a cocktail of experimental medications which must be personalised for the individual and are often poorly tolerated. Critically, with many patients, no therapeutic benefit is achieved. We sought to address this by using large-scale virtual screening to accelerate the development of a molecule which binds directly to KCNT1 to supress overactivity. We purchased a total of 71 compounds and using a combination of fluorescent thallium flux assays and patch clamp electrophysiology, identified a series of eight structurally diverse, novel inhibitors of the K_Na_1.1 channel with potency in the low micromolar range. These provide potential starting points for further development of drugs to treat *KCNT1*-associated epilepsy.

**Highlights:**

- We have discovered a range of structurally distinct inhibitors of the KCNT1 ion channel using large-scale virtual screening
- These compound exhibit selectivity for the KCNT1 channel over other related ion channels
- These compounds could provide starting points for new treatments for KCNT1 related Epilepsy.

**Graphical abstract:** 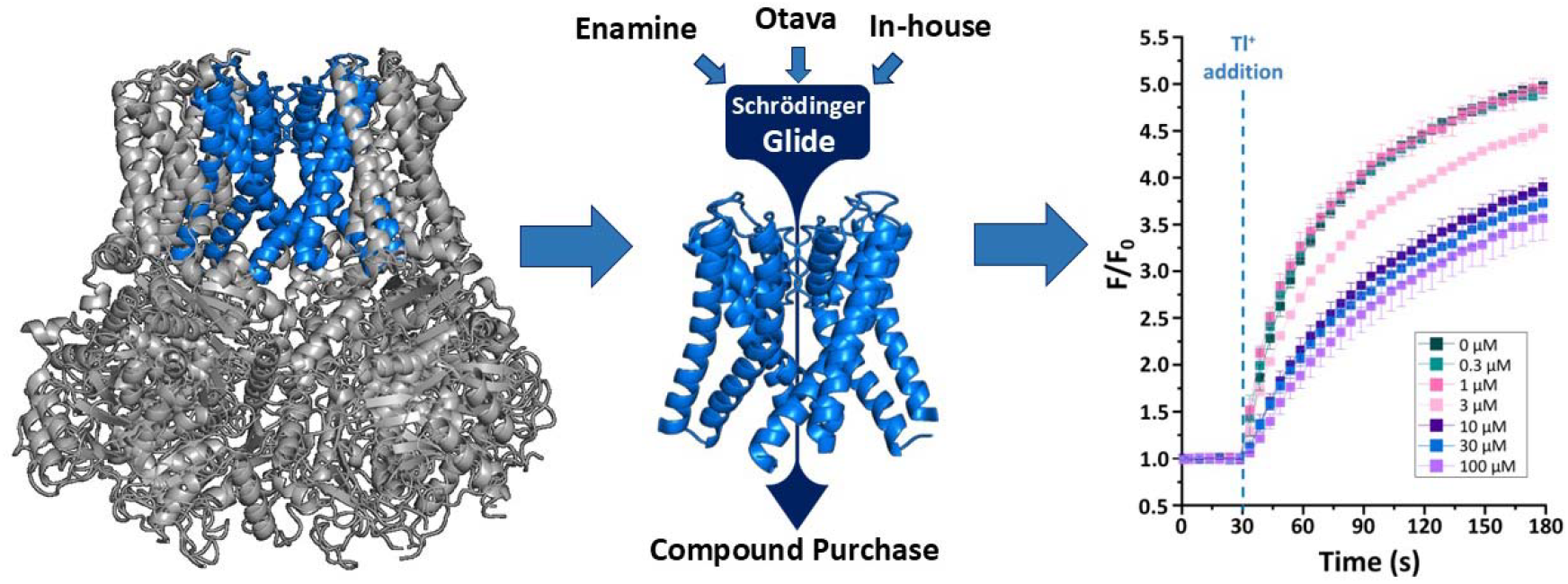

## Introduction

Potassium (K^+^) ion channels are among the most widely distributed group of ion channels and comprise a superfamily of structurally and functionally diverse proteins. Within this group, the SLO subfamily of four K^+^channels are among the most complex and exhibit unusually high conductance [1]. The *KCNT1* gene lies within this family and encodes the largest known potassium channel subunit, K_Na_1.1 (also known as SLACK, Slo2.2 or K_Ca_4.1) [2]. This subunit is composed of six transmembrane alpha helices, with a selectivity filter formed by a loop between the fifth and sixth helices, and two intracellular C-terminal regulation of conductance of potassium (RCK) domains [3, 4]. This subunit forms a homotetrameric sodium (Na^+^)-activated K^+^channel, but can also form heteromeric coassemblies with the closely related K_Na_1.2 subunit (encoded by *KCNT2*) where they colocalise in distinct regions of the central nervous system (CNS) [5, 6]. K_Na_1.1-containing channels are activated by millimolar concentrations of sodium (Na^+^) and contribute to Na^+^-dependent K^+^currents (*I*_*KNa*_). These channels play a role in generating the slow afterhyperpolarisation (AHP) following single [7, 8] or bursts of action potential firing [9], in addition to contributing to the resting membrane potential and intrinsic excitability of different cell types in the CNS [10, 11] and vascular smooth muscle [12].

Pathogenic gain-of-function (GoF) variants of the *KCNT1* gene are associated with rare and severe forms of drug-resistant epilepsy [13]. More than 50 reported missense variants have been implicated in a range of disorders, most commonly epilepsy of infancy with migrating focal seizures (EIMFS) [14] or autosomal dominant sleep-related hypermotor epilepsy (ADSHE) [15], with less common and more recently discovered phenotypes such as Ohtahara syndrome [16], West syndrome [17], and Lennox-Gastaut syndrome [18]. These conditions are often debilitating; patients present with recurrent seizures, with associated comorbidities including developmental delay or regression and intellectual disabilities [19, 20].

The refractoriness to conventional antiepileptic medications seen in these conditions has led to trials with alternative medications such as quinidine, a class I antiarrhythmic agent which inhibits K_Na_1.1 channels [21]. However, despite moderate improvements being observed in a small subset of patients, this treatment is largely ineffective [22-24] and can lead to severe cardiotoxicity due to quinidine’s lack of specificity and its high degree of potency at cardiac ion channels such as K_V_11.1 (hERG) [25]. Recently, a patient with the GoF *KCNT1* S937G mutation showed a drastic reduction in seizure activity following treatment with the antidepressant drug fluoxetine, which is also a K_Na_1.1 channel inhibitor [26]. However, whether this is a viable treatment strategy for the majority of patients within this heterogenous group of epilepsy phenotypes is yet to be seen from this n=1 study; as such, the discovery of further K_Na_1.1 channel inhibitors have potential for both the development of therapies and as tool compounds to interrogate functional properties of this channel in future studies.

To this end, efforts towards discovering small molecules for the specific pharmacological modulation of K_Na_1.1 potassium ion channels have seen an increase in recent years (Figure 1) [27-31]. Similarly, there have been recent advancements in the availability of high-resolution structural information available from cryo-electron microscopy (cryo-EM) structures of the K_Na_1.1 channel in different conformations [3, 4, 32]. We have previously shown that known K_Na_1.1 channel inhibitors typically bind to the inner pore vestibule [27], and the combination of this structural and functional understanding of the channel facilitates the use of a computer-aided approach for the discovery of novel channel modulators. Such as approach can drastically reduce the time and cost of drug development by precluding the requirement for the functional screening of large compound libraries [33].

**Figure 1:**
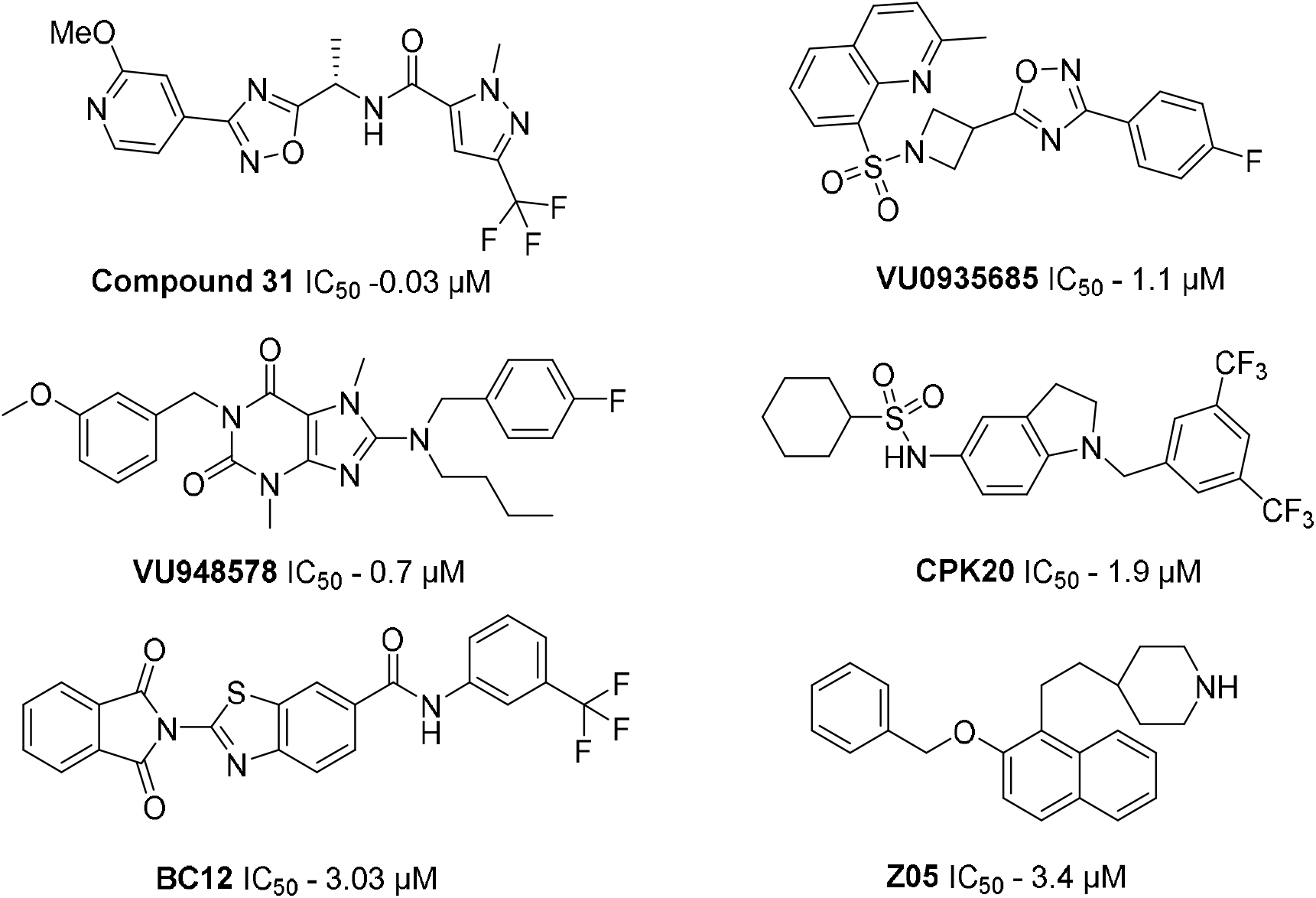
Structures of representative K_Na_1.1 potassium channel inhibitors and their IC_50_ values against the wild-type human K_Na_1.1 channel.

We aimed to build on our previous structure-based approach by significantly increasing the magnitude of the screen, using high-performance computing, from 10^5^compounds [27] to over 10^6^against the same target. We envisaged that this would increase the chemical diversity of our pool of inhibitors. In this study, we used a virtual high-throughput approach to screen a diverse library of commercially available compounds against the pore of the K_Na_1.1 cryo-electron microscopy structure [3]. Using a combination of fluorescent thallium flux assays and patch clamp electrophysiology, we identified a series of eight structurally diverse, novel inhibitors of the K_Na_1.1 channel with potency in the low micromolar range. Interestingly, new structural information which emerged following the discovery of these inhibitors [4] provided a further insight into the potential binding modes of these compounds, developing our capacity to expand our approach in future studies.

## Results and Discussion

### Structure-based virtual screening

In our previous work, we used a structure-based approach to identify the binding site of the known K_Na_1.1 potassium ion channel inhibitors, bepridil and quinidine, in the intracellular pore vestibule, proximal to the selectivity filter (Figure 2A). Informed by this, we virtually screened a library of 100,000 drug-like molecules against this site, and of the 17 compounds tested *in vitro*, six were identified as K_Na_1.1 inhibitors [27]. In the present study, we sought to determine whether we could scale this structure-based methodology to screen millions of compounds using high-throughput and high-performance computational tools. We performed virtual screening of different compound libraries against the same site in the chicken K_Na_1.1 cryo-electron microscopy structure in the putative active conformation (PDB 5U70), encompassing side chains from the selectivity filter and S6 transmembrane region which comprise the intracellular pore vestibule (Figure 2A). The amino acids contained in this region are highly conserved between the chicken and human proteins, with 96% sequence identity (Figure 2B). For this study we focused our efforts on commercially available libraries of compounds, beginning with our in-house library (MCCB) of approximately 25,000 compounds at the University of Leeds, in addition to the Enamine drug-like and Otava screening collection libraries, containing approximately 1.8 million and 270,000 compounds, respectively. For Enamine, the larger of the three libraries, we adopted a two-fold virtual screening approach wherein the compounds were initially docked using the fast docking tool, Fred (Openeye Scientific Software) using high-performance computing, after which the compounds with the top 10 % predicted binding scores from the Enamine library were docked along with the Otava and MCCB libraries using Glide (Schrödinger) in Standard Precision (SP) mode. Compounds from all libraries were ranked by their Glide SP docking score, and those with a docking score greater than -6 discarded. Any pan assay interference compounds (PAINS) [34] were removed, and from the top scoring compounds, 71 were selected based on structural diversity and obtained for further *in vitro* assessment. The workflow for this approach is shown in Figure 2C.

**Figure 2:**
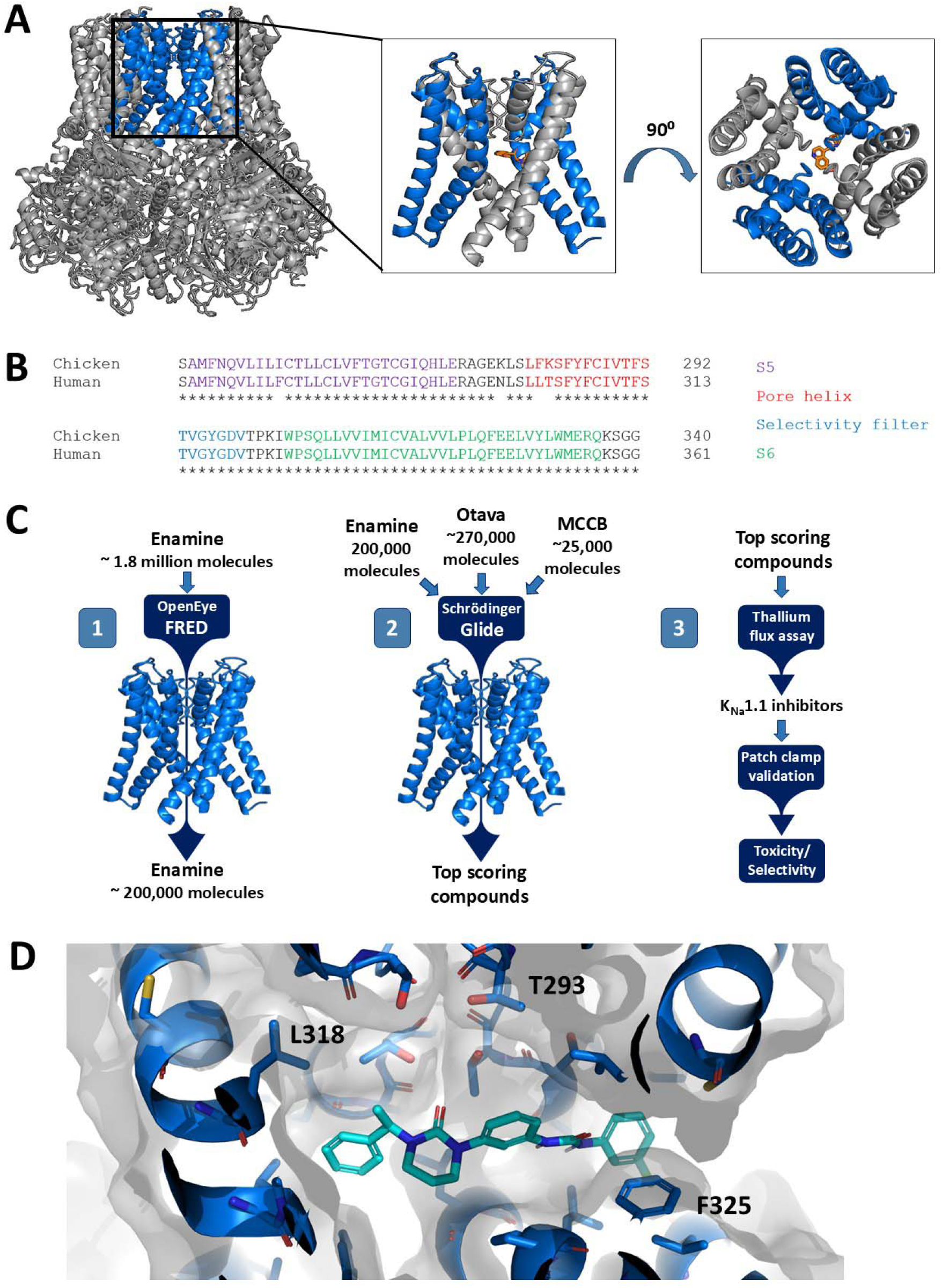
Virtual high throughput screening of chemical libraries to find inhibitors of K_Na_1.1 cryo-EM structure. A) The tetrameric K_Na_1.1 chicken cryo-EM structure in the open state (PDB 5U70) with the known inhibitor bepridil (orange) docked to a known binding site below the selectivity filter. Inset boxes show the 20 Å cubed region against which screening was carried out, shown parallel to the membrane (left) and from the extracellular side (right). B) Sequence alignment of the chicken and human K_Na_1.1 channel homologue sequences for the 20 Å cube used in virtual high throughput screens; conserved residues are indicated by an asterisk. C) Workflow for docking large scale libraries against the K_Na_1.1 structure, including selecting the top 10% predicted highest affinity binding molecules via an initial screen using FRED (OpenEye), then screening this smaller library using Glide (Schrödinger), after which the top scoring commercially available compounds were functionally screened for inhibition comparable to the positive control (100 µM bepridil). D) Predicted bound conformation of compound 1 in the open chicken K_Na_1.1 potassium ion channel structure (PDB 5U70). Protein shown as blue ribbons with key residues shown as blue sticks and compound 1 shown as cyan sticks.

### *In vitro* validation of selected compounds

To assess whether the compounds selected from the virtual high throughput screen act as K_Na_1.1 potassium ion channel modulators, we initially used a medium-throughput fluorescent thallium flux assay in HEK293 cells stably expressing a heteromer of the wild-type (WT) and GoF mutant Y796H K_Na_1.1 [15]. This heteromeric system was chosen to better represent the response of endogenous channels containing mutant K_Na_1.1 subunits *in vivo*; *KCNT1*-associated epilepsy patients are generally heterozygous, and channels formed by co-assemblies of the WT and mutant subunits display intermediate properties between homomeric channels [13]. Compounds were initially applied at 10 µM to determine their effects on thallium conductance through K_Na_1.1 channels and compared to cells treated with the known K_Na_1.1 inhibitor, 100 µM bepridil. Cells were loaded with thallium sensitive dye and incubated with or without compounds of interest, following which thallium sulphate was added after 30 seconds of background fluorescence readings. The slope of the resulting fluorescent reads following this addition was used to determine which compounds reduced thallium conductance to levels comparable to the bepridil control (Figure 3A). This equated to approximately 50% inhibition due to the presence of endogenous cation ion channels in HEK293 cells that also conduct thallium ions [35]. From the top compounds virtually screened from the MCCB, Enamine and Otava libraries, 19% showed levels of inhibition comparable to those displayed by the positive control. The potency of the K_Na_1.1 hits was initially evaluated using a thallium flux assay in cells pre-incubated in varying concentrations of the compound (Figure 3B and C), and any compounds determined to have an IC_50_ value <15 µM using this assay were taken forward for further pharmacological characterisation. The eight compounds that were selected for further analysis were all confirmed as inhibitors by patch clamp electrophysiology (Figure 3C; Table 1). A WST-1 Cell Viability Assay was used to examine compound toxicity in HEK293 cells. With the exception of compound 3, none affected cell viability at concentrations up to 30 µM (Figure 4).

**Table 1:**
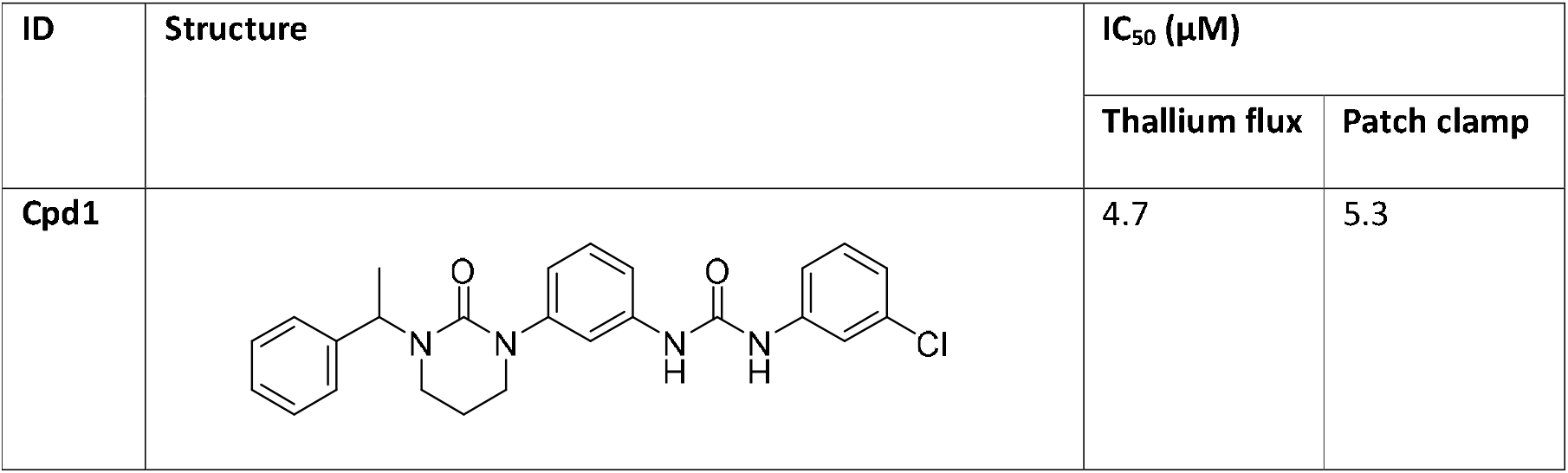
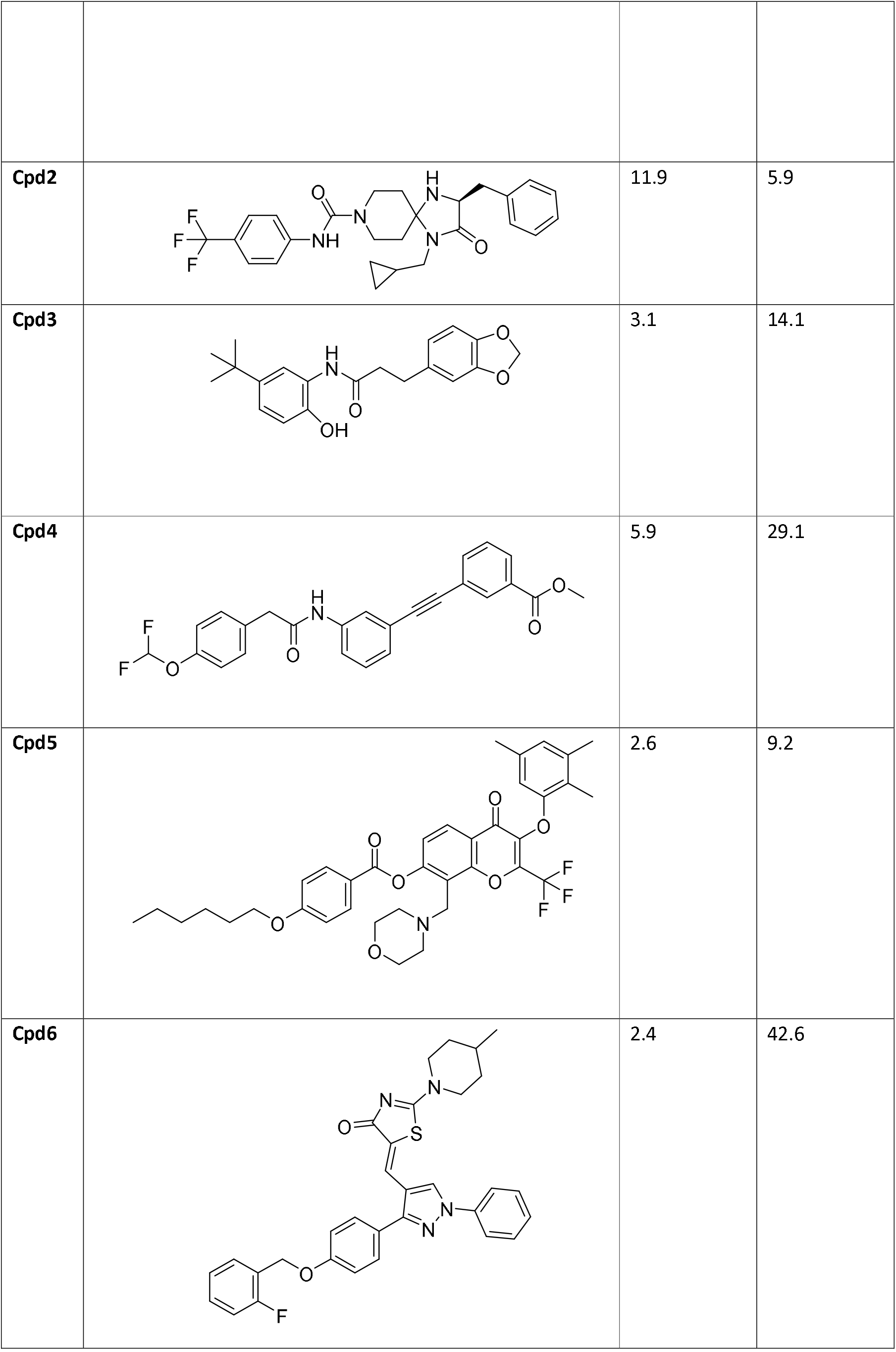
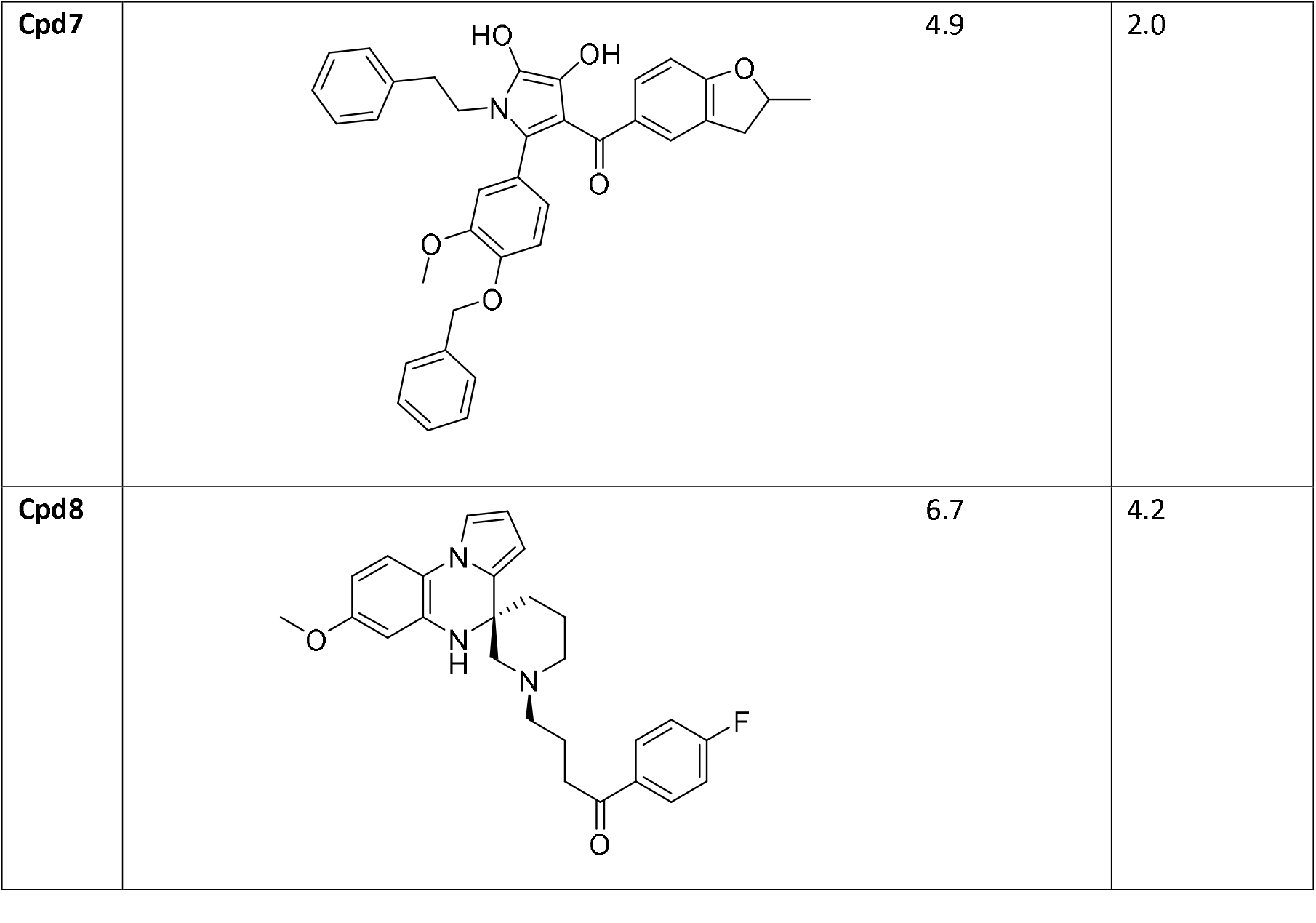
Selected compounds and their potencies (mean IC_50_ values) in inhibiting K_Na_1.1 channels measured using thallium fluorescence and validated by patch clamp electrophysiology.

| ID | Structure | IC<sub>50</sub> (μM) |  |
| --- | --- | --- | --- |
|  |  | Thallium flux | Patch clamp |
| Cpd1 |  | 4.7 | 5.3 |
| Cpd2 |  | 11.9 | 5.9 |
| Cpd3 |  | 3.1 | 14.1 |
| Cpd4 |  | 5.9 | 29.1 |
| Cpd5 |  | 2.6 | 9.2 |
| Cpd6 |  | 2.4 | 42.6 |
| Cpd7 |  | 4.9 | 2.0 |
| Cpd8 |  | 6.7 | 4.2 |

**Figure 3:**
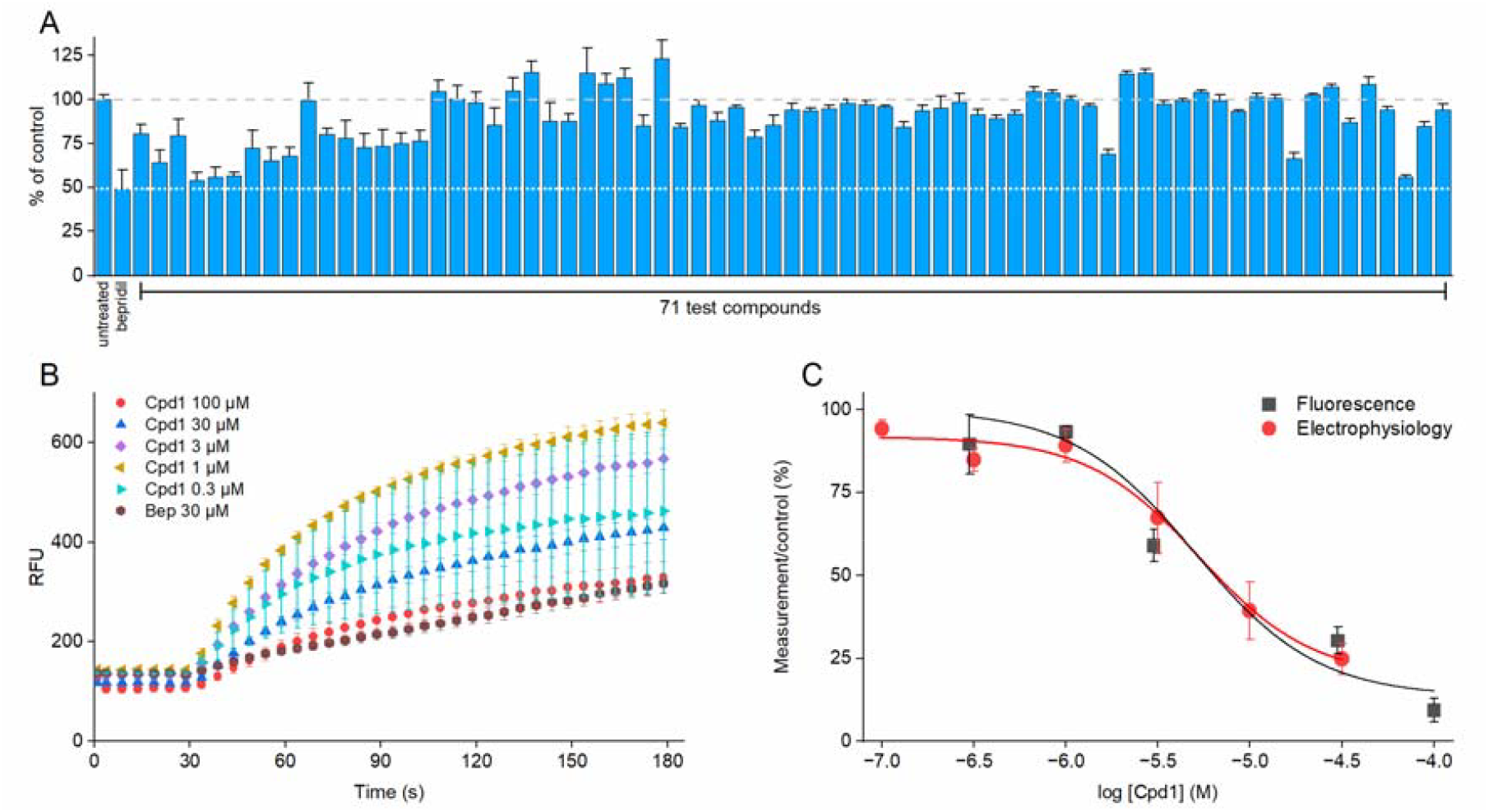
*In vitro* validation of top scoring compounds identified through virtual high throughput screening. A) Thallium flux measured from stable HEK293 K_Na_1.1 WT/Y796H cells incubated with test compounds at 10 µM. The grey dashed line indicates the control response in the absence of inhibiter and the white dotted line indicates the inhibition by the positive control, 100 µM bepridil. B) Representative mean thallium flux data showing thallium-induced fluorescence in response to cell incubation with varying concentrations of compound 1 C) Concentration-inhibition curves of compound 1 assessed using thallium fluorescence and patch clamp electrophysiology.

**Figure 4:**
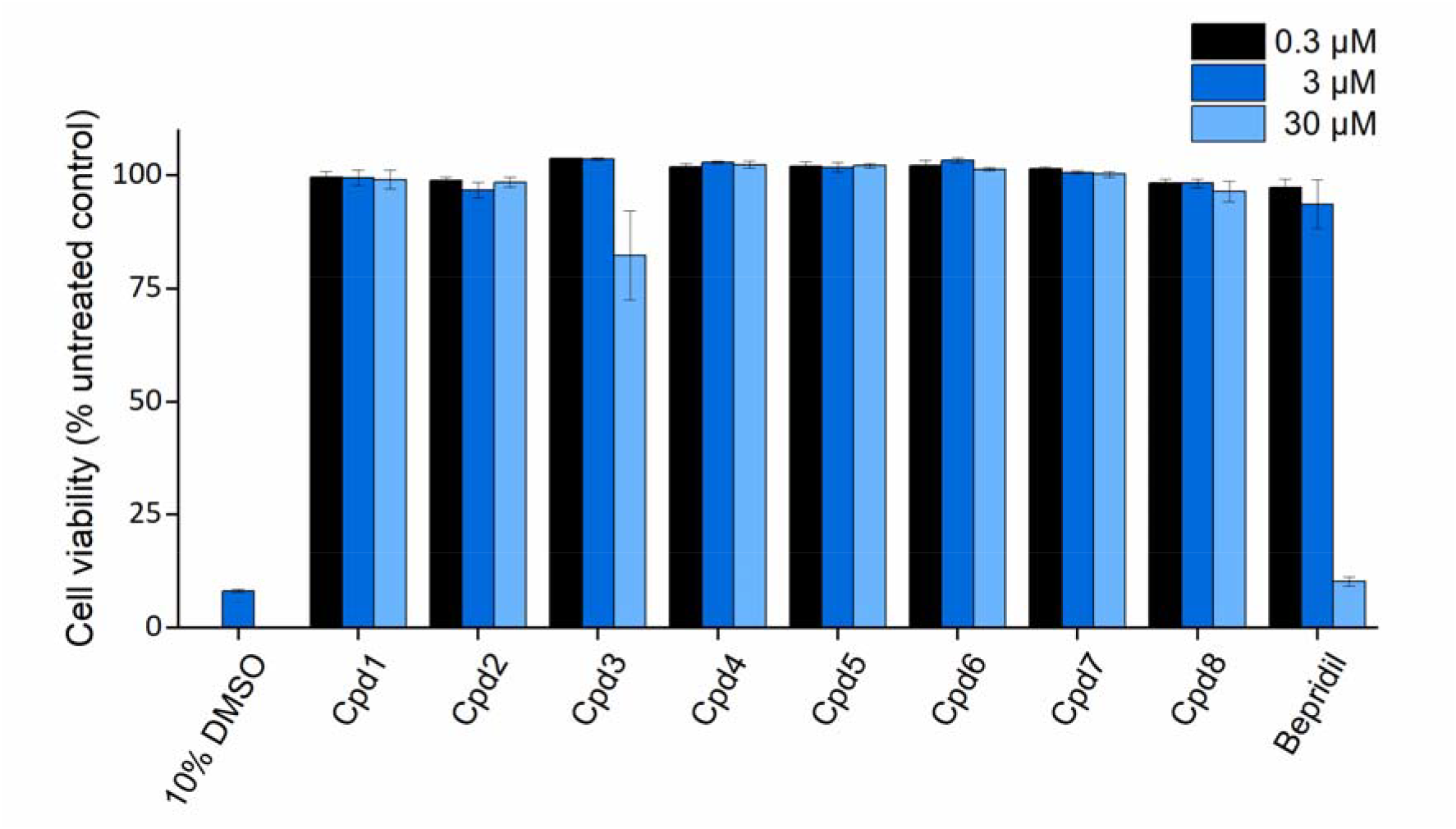
Summary of percentage of cell viability of HEK293 cells treated for 24 hours with the indicated concentration of compound, as determined by WST-1 assay, with 10 % DMSO as a positive control.

### K_Na_1.1 inhibitor selectivity

The selectivity profile of quinidine as a treatment for KCNT1-associated has been a significant barrier to reaching therapeutic doses without significant adverse effects. Further characterization of compounds 1-8 included an assessment of potential secondary activity across a panel of seven other potassium channels, including channels encoded by other members of the *Slo* gene family, at 10 μM compound concentration using the thallium flux assay in HEK293 cells stably expressing channels of interest (Table 2). Pleasingly, most compounds showed a relatively clean selectivity profile, with only compounds 1 and 3 showing greater than 20% channel inhibition for K_V_4.2. Blockade of hERG potassium channels is a major cause for pro-arrhythmic effects associated with QT prolongation and torsade-des-pointes in humans. Using patch clamp, it was determined that a number of these compounds (2,3 and 6) inhibit the hERG channel by over 50% at 10 µM (Figure 5). Conversely, several of these compounds, for example compound 7 and 8 exhibit very little inhibition of the channel, demonstrating the value of the *in-silico* approach to identify diverse novel channel modulators.

**Figure 5:**
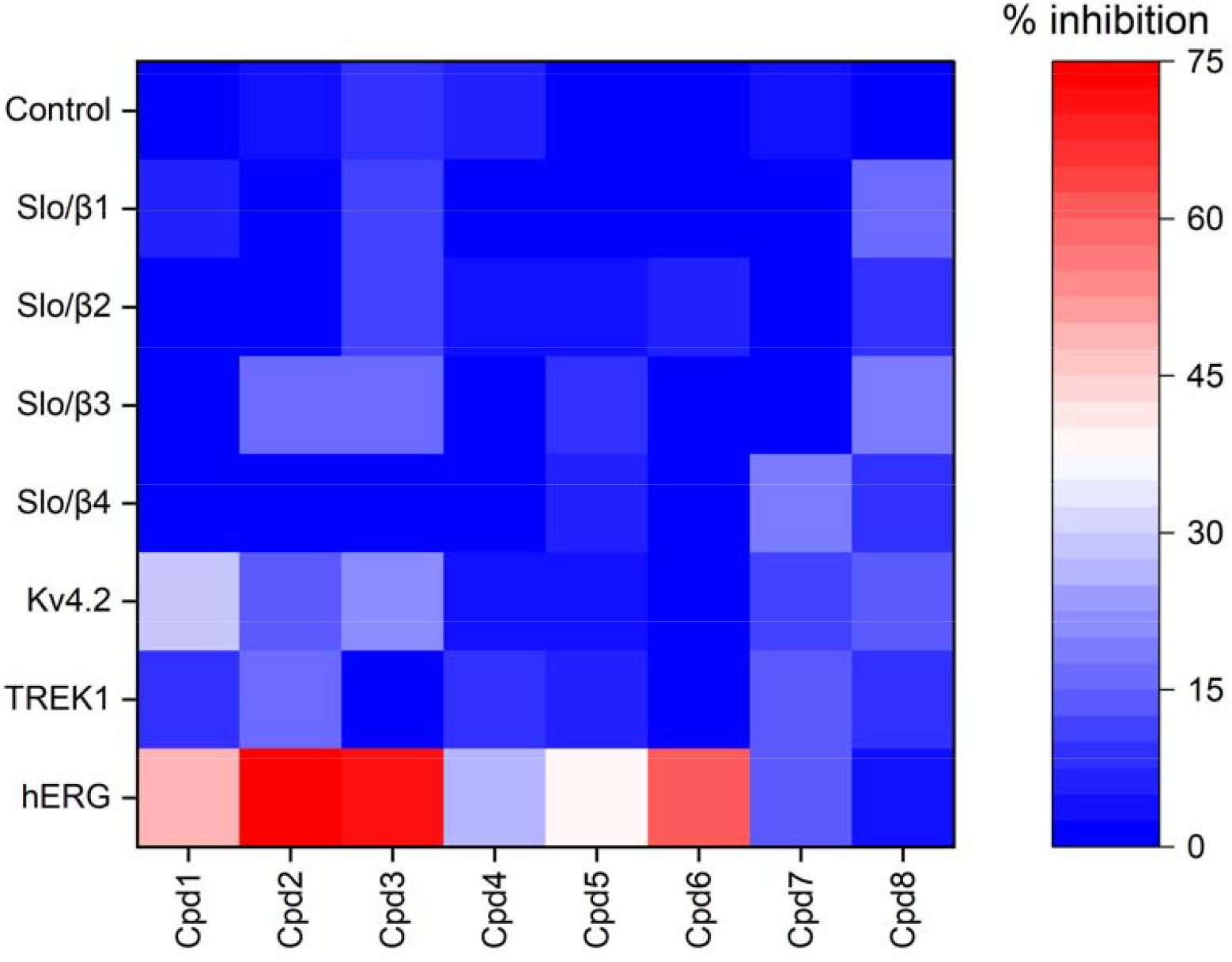
Heatmap indicating mean % inhibition of control HEK293 cells or those expressing potassium channels as indicated by 10 µM of Compounds 1 - 8. Data were obtained using thallium flux assay except for hERG, which was analysed using patch clamp electrophysiology.

### Predicted binding mode in the human K_Na_*1*.1 structure

The virtual high throughput screening was carried out using the cryo-EM structure of the pore region of the chicken K_Na_1.1 channel [3] as an expansion of our previous research [27]. The chicken and human proteins are highly homologous, with 96% sequence identity within the region targeted in our virtual high throughput screening (Figure 2A). The sequence of the inner pore vestibule is conserved between species (Figure 2B), meaning that the chicken K_Na_1.1 structures are likely to be a reliable model for the interaction of inhibitors with the human K_Na_1.1 channels. More recently, cryo-EM structures were published of the human K_Na_1.1 channel [4]. This series of structures showed the human K_Na_1.1 ion channel in open, closed and inhibitor-bound states, at resolutions ranging between 2.6–3.2 Å. The inhibitor-bound structure elucidated for the first time, the binding mode of a potent K_Na_1.1 channel inhibitor [29]. Arguably the most pivotal revelation to emerge from this series of structures, from an inhibitor binding perspective, was the discovery of a new binding pocket in the ligand-bound state with an inhibitor wedged into a pocket formed by pore helix and S6 helix, blocking the pore. The ligand-bound human structure showed movement of the Phe312 side chain into a previously unseen pose, exposing a new binding pocket that could accommodate part of the inhibitor. Future *in silico* studies can utilise these structures to further improve the screening cascade for inhibitor identification.

## Conclusions

This study has identified a number of novel, structurally diverse, and synthetically tractable inhibitors of the human K_Na_1.1 ion channel using a combination of *in silico* large-scale virtual high-throughput screening, *in vitro* thallium flux assays and whole cell patch-clamping. This large-scale approach has led to the development of more structurally diverse and synthetically tractable compounds compared to our previous smaller screen [27]. We have shown it is possible to achieve selectivity for the K_Na_1.1 channel versus other potassium channels, including hERG, which is crucial for further development of lead candidates. Due to the lack of commercially available analogues of these compounds, we are currently unable to develop the structure-activity relationship profile of these inhibitors to understand their requirements for binding. This has also limited the optimisation of the pharmacokinetic properties of these inhibitors, many of which suffer from poor aqueous solubility and low metabolic stability. The recent publication of an inhibitor-bound cryo-EM structure of human K_Na_1.1 provides an enhanced scope to use the approach discussed here for the future identification and optimisation of inhibitors to treat KCNT1-related neurological disorders.

## Materials and Methods

### Molecular biology and cell culture

The full-length wild-type (WT) K_Na_1.1 cDNA in pcDNA6-V5/His6 vector (Invitrogen) is described previously (Cole et al, 2020). Using this, a concatemeric construct was generated in pcDNA6 so that a second K_Na_1.1 subunit containing the known Y796H GoF variant was fused to the C-terminus of the wild type K_Na_1.1 with a T2A “self-cleaving” linker sequence [36]. This construct was transfected into human embryonic kidney (HEK) 293 cells (transfection and culture details provided below) and cultured in medium containing 5 μg/mL blasticidin for selection of stably transfected clones. Blasticidin-resistant colonies of HEK cells were isolated, expanded, and assessed for stable WT/Y796H K_Na_1.1 currents by whole cell patch clamp recording (see electrophysiology section below).

Human embryonic kidney (HEK) 293 cells, either stably expressing or transiently transfected with the ion channel of interest, were cultured at 37 °C in air containing 5% CO_2_ in Dulbecco’s modified Eagle medium (DMEM with GlutaMax, Invitrogen), supplemented with 10 % foetal bovine serum, 50 U/mL penicillin, and 0.05 mg/mL streptomycin. When required, cells were transiently transfected using the TransIT-X2 reagent (Cambridge Bioscience) according to the supplier’s specifications.

### Compound library generation

Compound libraries from Otava (www.otavachemicals.com) and Enamine (https://enamine.net/) were downloaded in SMILES format. Three-dimensional versions of these libraries were produced using OpenEye OMEGA conformer generation software (OpenEye OMEGA Application 2022.1.1, OpenEye, Cadence Molecular Sciences, Santa Fe, NM. http://www.eyesopen.com.) [37]. Due to the large size of these libraries, only the lowest energy conformer was produced for each compound.

### Virtual screening

To identify candidate compounds, high-throughput virtual screening was carried out using the ARC4 advanced research computing cluster at the University of Leeds. Cryo-electron microscopy structures of the chicken K_Na_1.1 potassium ion channel (PDB 5U76 or 5U70 [3] for the inactive and active states, respectively), were prepared using the Maestro protein preparation wizard (Schrödinger Release 2020-3, Schrödinger, LLC, New York, NY, 2024). Large libraries of commercially available compounds from Enamine and Otava, as well as an in-house library, were initially screened using the FRED docking algorithm (OpenEye FRED Application 2022.1.1, OpenEye, Cadence Molecular Sciences, Santa Fe, NM. http://www.eyesopen.com) [38] using the Chemgauss4 scoring function. Compound libraries were docked to a region of the pore domain comprising a 20 Å cube encompassing side chains from the selectivity filter and S6 transmembrane segment that line the intracellular pore vestibule (Figure 1A). The compounds were ranked according to their Chemgauss4 docking score and the 10% top-scoring compounds subsequently rescored using Schrödinger Glide [39] in Standard Precision (SP) mode. Approximately 500 compounds were visually inspected using Maestro to identify predicted specific binding interactions with the protein (Figure 2D) and disregard compounds with pan-assay interference structures [34]. Structurally diverse compounds with the best docking scores were purchased and tested *in vitro*.

### Cell viability

Cell viability was determined using the WST-1 assay according to the manufacturer’s instructions. HEK293 cells were seeded into a 96 well plate at a concentration of 5 x 10^4^ cells per well in 100 µL of DMEM culture medium and incubated at 37°C with 5% CO_2_ overnight. Compounds of interest or positive control, consisting of 10% DMSO, were subsequently added to wells in triplicate and the cells incubated for a further 24 hours, after which 10 µL of WST-1 reagent (Merck, 5015944001) was added to each well. Cells were incubated at 37°C with 5% CO_2_ for 4 hours, then thoroughly shaken for 1 minute before reading the absorbance of the plate 450 nm, reference wavelength 650 nm. Cell viability was calculated as a percentage of the absorbance read from untreated cells.

### Thallium flux assay

All chemicals were sourced from Sigma-Aldrich (Gillingham, U.K.) unless otherwise stated. HEK293 cells were plated into a clear-bottomed, poly-D-lysine coated 96-well plate (ThermoFisher Scientific) at a density of 5 x 10^5^ cells per well in 100 µL of DMEM culture medium and incubated at 37°C with 5% CO_2_ overnight. Cells were then washed once with chloride-free Hank’s balanced salt solution (HBSS) (140 mM sodium gluconate, 5 mM potassium gluconate, 1 mM calcium gluconate, 0.9 mM magnesium sulphate heptahydrate, 0.3 mM sodium phosphate dibasic dihydrate, 0.4 mM potassium phosphate monobasic, 6 mM D-Glucose, 4 mM sodium bicarbonate, pH 7.3) and loaded with 90 µL of assay buffer consisting of 1.25 µg/mL Thallos AM dye (ION Biosciences), 0.1 % Pluronic™ F-127 (ThermoFisher Scientific) and extracellular fluorescence quenchers (5 mM tartrazine and 10 mM amaranth). 10 µL of test compound were diluted from a 10 mM DMSO stock with chloride-free HBSS and quenchers and the plate incubated at 37°C for 1 hour. For initial validation, compounds were applied at a final concentration of 10 µM, followed by a range of concentrations for ‘hit’ molecules for concentration-inhibition analysis. Following incubation, Thallos fluorescence was read every 5 seconds using a FlexStation 3 Microplate reader (Molecular Devices) (excitation 490 m, emission 515 nm), wherein baseline fluorescence was measured for 30 seconds prior to the addition of thallium sulphate solution in HBSS with quenchers to a final concentration of 2.5 mM, after which fluorescence was measured for a further 150 seconds. The slope of the first ten measurements versus time, following thallium sulphate addition, was used as a measure of channel activity, and the slopes from the untreated wells were used as control.

### Manual patch-clamp electrophysiology

Membrane currents were measured at room temperature using patch-clamp recording in the whole-cell configuration. An EPC-10 amplifier (HEKA Electronics, Lambrecht, Germany) was used, with >65 % series resistance compensation (where appropriate), 2.9 kHz low-pass filtering, and 10 kHz digitisation as described previously [27]. HEK293 cells were seeded onto 10 mm glass cover slips a day prior to use and placed into a recording chamber connected to a gravity perfusion system. Patch pipettes with a resistance of 1.5-3.0 MΩ in experimental solution were pulled from thin-walled borosilicate glass (Harvard Apparatus Ltd, Edenbridge, Kent, UK) and polished. The extracellular recording solution for recording K_Na_1.1 potassium currents contained 140 mM NaCl, 1 mM CaCl_2_, 5 mM KCl, 29 mM Glucose, 10 mM HEPES and 1 mM MgCl_2_, pH 7.4 with NaOH, and the pipette solution contained 100 mM K-Gluconate, 30 mM KCl, 10 mM Na-Gluconate, 29 mM Glucose, 5 mM EGTA and 10 mM HEPES, pH 7.3 with KOH. To assess the current-voltage relationship, cells were kept at a holding potential of -80 mV and 400 ms pulses were applied at 10 mV increments between -100 and 80 mV. Inhibition by hit compounds from the thallium flux screen was assessed by the application of compounds via gravity perfusion, followed by concentration-inhibition analysis: G/G_C_ = (1 + ([B]/IC _50_)^H^)^-1^+ c, where G is the conductance measured as the slope of the current evoked by the voltage ramp in the presence of the inhibitor, G_C_ is the control conductance in the absence of inhibitor, [B] is the concentration of the inhibitor, IC_50_ the concentration of inhibitor that yields 50% inhibition, H the slope factor, and c the residual current.

The extracellular recording solution when recording cells expressing the hERG potassium ion channel contained 137 mM NaCl, 1.8 mM CaCl_2_, 4 mM KCl, 10 mM Glucose, 10 mM HEPES, 1 mM MgCl_2_, pH 7.3 with NaOH and the pipette solution contained 140 mM KCl, 10 mM EGTA, 10 mM HEPES, 2 mM MgCl_2_, 5 mM Mg_2_ATP, pH 7.2 with KOH. To assess channel inhibition, the cells were held at -80 mV and subsequent 4 second pulses 40 mV and -50 mV applied. Compounds were applied by gravity perfusion at the indicated concentrations and the degree of inhibition determined from the tail current amplitude. Data analysis was conducted using Fitmaster (HEKA Electronics, Lambrecht, Germany), Microsoft Excel, and OriginPro 9.65 (OriginLab Corporation, Northampton, MA, USA).

## Acknowledgements

This work was undertaken on ARC4, part of the High-Performance Computing facilities at the University of Leeds, UK.

## Author contributions

Emily A. Casely: Methodology, Investigation, Formal analysis, Visualization, Writing - Original Draft; Katie J. Simmons: Methodology, Conceptualization, Supervision, Writing - Original Draft, Funding acquisition; Alex Flynn: Methodology, Investigation; Bethan A. Cole: Methodology, Resources; Stephen P. Muench: Conceptualization, Writing - Review & Editing, Funding acquisition; Jonathan D. Lippiat: Conceptualization, Resources, Visualization, Supervision, Writing - Original Draft, Funding acquisition, Project administration.

## Data Availability

Non-confidential research data will be available from the corresponding author upon request.

